# Disruption of redox balance in glutaminolytic triple negative breast cancer by inhibition of glutamate export and glutaminase

**DOI:** 10.1101/2023.11.19.567663

**Authors:** Hoon Choi, Mamta Gupta, Christopher Hensley, Hsiaoju Lee, Yu-Ting Lu, Austin Pantel, David Mankoff, Rong Zhou

## Abstract

In triple-negative breast cancer (TNBC) that relies on catabolism of amino acid glutamine, glutaminase (GLS) converts glutamine to glutamate, which facilitates glutathione synthesis by mediating the enrichment of intracellular cystine via xCT antiporter activity. To overcome chemo resistant TNBC, we have tested a strategy of disrupting cellular redox balance by inhibition of GLS and xCT by CB839 and Erastin, respectively. Key findings of our study include: 1. Dual metabolic inhibition (CB839+Erastin) led to significant increases of cellular superoxide level in both parent and chemo resistant TNBC cells, but superoxide level was distinctly lower in resistant cells. 2. Dual metabolic inhibition combined with doxorubicin or cisplatin induced significant apoptosis in TNBC cells and is associated with high degrees of GSH depletion. *In vivo*, dual metabolic inhibition plus cisplatin led to significant growth delay of chemo resistant human TNBC xenografts. 3. Ferroptosis is induced by doxorubicin (DOX) but not by cisplatin or paclitaxel. Addition of dual metabolic inhibition to DOX chemotherapy significantly enhanced ferroptotic cell death. 4. Significant changes in cellular metabolites concentration preceded transcriptome changes revealed by single cell RNA sequencing, underscoring the potential of capturing early changes in metabolites as pharmacodynamic markers of metabolic inhibitors. Here we demonstrated that 4-(3-[^18^F]fluoropropyl)-L-glutamic acid ([^18^F]FSPG) PET detected xCT blockade by Erastin or its analog in mice bearing human TNBC xenografts. In summary, our study provides compelling evidence for the therapeutic benefit and feasibility of non-invasive monitoring of dual metabolic blockade as a translational strategy to sensitize chemo resistant TNBC to cytotoxic chemotherapy.

## Introduction

Triple negative breast cancer (TNBC) is a complex and aggressive subtype of breast cancer that poses unique challenges in treatment due to the absence of estrogen receptors (ER), progesterone receptors (PR), and human epidermal growth factor receptor 2 (HER2). Unlike other subtypes that benefit from targeted therapies, such as endocrine treatment for ER/PR-positive or HER2-targeted therapy for HER2-expressing breast cancers, TNBC lacks subtype-specific treatment options. Systemic therapy in the form of chemotherapy remains an important component of treatment for nearly all TNBC patients as adjuvant or neoadjuvant therapy to nonmetastatic disease and as primary therapy to metastatic TNBC. Despite the initial positive responses to chemotherapy, resistance develops frequently, which leading to cancer relapse or progression, highlighting the need for innovative and translational strategies to overcome chemotherapy resistance (1,2).

Utilizing glutamine as an essential nutrient for survival and growth has been recognized as a metabolic vulnerability of aggressive cancers including TNBC (3-7). To target glutaminolysis pathway (purple arrow in **SI Figure 1**), a highly specific and potent inhibitor of kidney-type glutaminase (GLS), CB839 (Telaglenastat) has been developed (5) that blocks the conversion of glutamine (Gln) to glutamate (Glu), the first and rate-limiting step of glutaminolysis. Clinical trials of CB839 have demonstrated an excellent safety profile, yet the anti-cancer efficacy observed has been variable (8-10).

Emerging research has unveiled that Glu, at the intersection of glutaminolysis and *de novo* glutathione (GSH) synthesis pathway (green arrows in SI Figure 1) may play an important role in maintaining cellular redox homeostasis. Reactive oxygen species (ROS) are produced from physiological processes of cells including OXPHOS in mitochondria, NADPH oxidase activity, growth factor receptor engagement, and exposure to xenobiotics or radiation (11,12). Due to aberrations in metabolic and growth signal regulation, cancer cells exhibit elevated ROS level than normal cells, hence having a greater demand for cellular antioxidants because unmitigated ROS lead to disruption of cellular redox balance, triggering cell death via apoptosis (13) and/or ferroptosis mediated by lipid peroxidation (14-16). GSH synthesis pathway is essential even at the initiation of TNBC (17,18), consistent with GSH being as an essential cellular antioxidant to maintain redox homeostasis at all stages of cancer evolution including initiation, progression, metastasizing, and responses to treatment (19).

Cellular Glu contributes to GSH synthesis in two main aspects: first, via xCT (SLC7A11) antiport activity (20), Glu facilitates the enrichment of intracellular cystine, a rate limiting substrate for GSH synthesis (21); second, Glu is directly incorporated into GSH molecule, a L-γ-glutamyl-L-cysteinyl-glycine tripeptide. Expression of xCT is prevalent in human TNBC (22). However, xCT inhibition by small molecules such as Erastin or its analogs showed limited efficacy in clinical trials (23). As demonstrated in lung cancer model, Glu utilized for GSH synthesis would limit its availability for the TCA cycle, especially when cellular Glu pool is reduced by GLS blockade via CB839 (24). The mutual engagement of glutaminolysis and GSH synthesis with Glu being the nexus of the two pathways has motivated us to examine a dual metabolic inhibition approach (SI Figure 1). In studies presented herein, we tested the hypothesis that pharmacological blockade of xCT and GLS deplete cellular GSH, leading to unbalanced redox state that drives apoptosis and/or ferroptosis in TNBC cells. We explored this approach for overcoming chemo resistant TNBC by *in vitro* and *in vivo* studies. The outcomes of these investigations suggest that this translational strategy holds significant promise in sensitizing resistant TNBC to chemotherapy.

## Materials and Methods

Human TNBC cell line (HCC1806) was purchased from ATCC (catalog CRL-2335). HCC1806R is a paclitaxel resistant line derived from HCC1806 (parent) that was exposed to incremental concentration of paclitaxel as previously described (25). The cell lines were authenticated using the short-tandem-repeat DNA profiling method. GLS inhibitor (GLSi) CB839 (Calithera Biosciences, Palo Alto, CA) was formulated in dimethyl sulfoxide for cell studies or in vehicle solution as described earlier (5) for in vivo administration. The following chemicals were purchased from Cayman Chemical (Ann Arbor, MI): xCT inhibitor (xCTi) Erastin (ERA) and its analog, Imidazole ketone Erastin (IKE); from ThermoFisher Scientific: Annexin V-FITC, TO-PRO-3, dihydroethidium (DHE, catalog D11347), C11-Bodipy (catalog D3861), and buffer (catalog 00-4222-26) for Fluorescence-activated cell sorting (FACS); from Sigma or Millipore Sigma: Accumax (catalog A7089), glutamate (catalog D5030), doxorubicin (DOX), cisplatin (CIS) and paclitaxel (PTX) in pharmaceutical grade.

### In vitro and ex vivo studies

TNBC cells were cultured in RPMI1640 media (catalog MT10-040-CM, Corning, NY) supplemented with 10% FBS (catalog MT35-010-CV, Corning, NY). No antibiotics were used in the culture.

#### Estimation of superoxide and lipid peroxidation level (ferroptosis) in cancer cells

HCC1806 and HCC1806R cells were initially seeded in a 12-well plate and allowed to culture overnight in culture media (RPMI1640+10% FBS). Subsequently, the media was replaced with culture media containing specific inhibitors and/or chemotherapy drugs: CB839 (1 μM), ERA (3 μM), DOX (0.2 μM), and the cells were then cultured for an additional 24 hours followed by incubation with 10 μM DHE (probe for superoxide) or 2 μM C11-Bodipy (probe for lipid peroxidation / ferroptosis) as final concentration at 37°C for additional 30 minutes. For combination treatments involving two or three drugs, the concentrations of each drug remained the same as used for the single-agent treatment. Following three washes with PBS, the cells were detached using Accumax diluted with FACS buffer. Fluorescent intensities of the oxidized DHE and oxidized C11-Bodipy, were measured using FACS (BD Bioscience), and the data were analyzed using FlowJo software. Specified fluorophore (bandpass filter wavelength in nm/width)/cell number were as follows: oxidized C11-Bodipy (530/30 nm)/10,000 cells and oxidized DHE (610/20 nm)/10,000 cells.

#### Estimation of early apoptosis by FACS/cell sorting

Early apoptotic cells and dead cells were assessed by Annexin V-FITC and TO-PRO-3 staining, respectively followed by FACS and cell sorting. Cells were cultured in media (RPMI1640 including 10% FBS) with specified drug or their combinations, including CB839, Erastin, doxorubicin (DOX) or cisplatin (CIS) for 24 hours with concentrations specified in Figure 1 caption for HCC1806 and HCC1806R cells, respectively. Following the incubation period, the cells were collected and washed 3 times with PBS. The collected cells were resuspended in 100 μL of HEPES buffer solution (10 mM HEPES, pH 7.4). Annexin V-FITC (5 μL,) and TO-PRO-3 (20 μL) were added to the cell suspension and incubated at room temperature for 15 minutes. The cell populations were assessed by FACS on a BD LSRII flow cytometer (BD Bioscience), utilizing fluorophore (bandpass filter wavelength in nm/width)/cell number as described: Annexin V-FITC (530/30 nm)/10,000 cells, and TO-PRO-3 (660/20 nm)/10,000 cells. The acquired FACS data were analyzed using FlowJo software (BD Bioscience) to determine the extent of early apoptosis under various treatment conditions.

**Figure 1.**
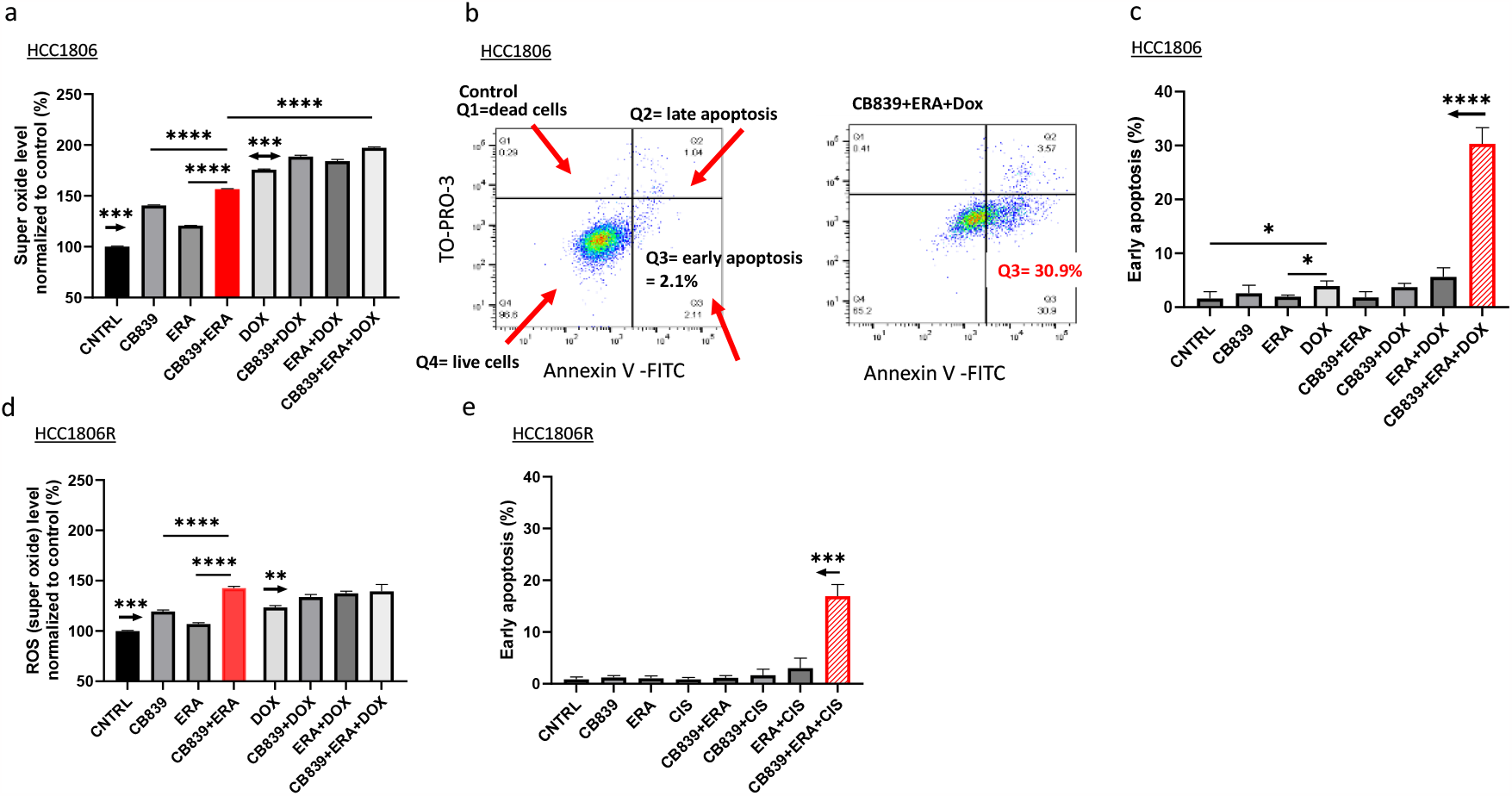
Dual metabolic blockade and chemotherapy drugs increased cellular superoxide level and mediated apoptosis in chemo naive and resistant TNBC cells. **a**: Super oxide level (normalized to untreated controls set at 100%) in cells exposed to CB839, ERA, DOX or their combinations (N=4 replicates for each treatment). In a, the arrow above CNTRL indicates *P*<0.001 comparing with each treatment in the panel, while the double arrow above the DOX indicates *P*<0.001 comparing with each treatment on the left or right of the DOX. **b:** Detection of early apoptosis by FACS using dual probes (additional treatments were described in SI Figure 2). **c:** Percentage of cells undergoing early apoptosis after the treatments specified in **a** and SI Figure 2 (N=3 for each treatment). In c, the arrow indicates *P*<0.0001 comparing CB839+ERA+DOX with CNTRL or any other treatment in the panel. **d:** Super oxide level in chemo-resistant cells exposed to the same treatments as in a (N=4 replicates for each treatment). In d, the arrow above CNTRL indicates *P*<0.001 comparing with any treatment in the panel, while the arrow above DOX indicates *P*<0.01 comparing with any DOX+ treatment on the right. **e:** Percent of early apoptosis in chemo-resistant cells exposed to CB839, ERA, CIS or their combinations (N=3 replicates for each treatment). The arrow indicates *P*<0.001 comparing CB839+ERA+CIS with CNTRL or any other treatment in panel e. Mean and SD were presented for all panels except for b. For early apoptosis measurement (c, e), HCC1806 cells were incubated with ERA (3μM), CB839 (1μM), Dox (0.2μM), or their combinations for 24 h, while HCC1806R cells were incubated with ERA (6μM), CB839 (2μM), CIS (10μM), or their combinations for 24 h.

#### Estimation of intra and extracellular Glu concentrations

One million cells were seeded per well in a 10 cm cell culture dish and incubated in culture media without glutamate (Glu) but containing glutamine (0.584g/l), glucose (1g/l), and NaHCO3 (3.7g/l) with 10% FBS for 24 hours at 37°C. The following day, the cells were treated with CB839 (6 μM), ERA (18 μM), or DOX (1.2 μM) in the same media without FBS for 6 hours. After the incubation, both the media and cells were collected. The cells were collected by scraping in 1 ml of ice-cold PBS. Subsequently, both the media and cells were lyophilized using Labconco™ FreeZone™ 4.5L 

~~~
(Fisher Scientific)
~~~

 and kept in -80°C freezer. To determine the Glu content in the samples, the lyophilized powder was reconstituted with deionized (DI) water and processed using the Sigma Glutamate Assay Kit (catalog no: MAK330, Sigma Aldrich) while the protein content was measured using the Pierce™ BCA Protein Assay Kit (catalog no: 23225, ThermoFisher Scientific), following manufacturers’ instructions.

#### Estimation of Glutathione (GSH) level in cancer cells

200K cells were seeded per well in 12-well plate and cultured in RPMI1640 media supplemented with 10% FBS for 24 hours at 37°C with 5% CO2. The next day, HCC1806 cells were treated with culture media containing the following drugs or their combination: CB839 (1 μM), ERA (3 μM), or DOX (0.2 μM) at 37°C for 24 hours. After the treatment, both the control (not exposed to any of the drugs) and treated cells were detached using trypsin and centrifuged at 700 x g for 5 minutes at 4°C. The supernatant was removed, and the cell pellet was resuspended in 0.5 mL of ice-cold PBS. The resuspended cells were then transferred to a 1.5 mL microcentrifuge tube and centrifuged at 700 x g for 5 minutes at 4°C. The supernatant was again removed, and the cells were lysed in 80 μL of ice-cold buffer from Glutathione Assay Kit (catalog no: ab239709, Abcam). The lysed cells were kept on ice for 10 minutes followed by mixing thoroughly with 20 μL of 5% SSA (sulfosalicylic acid) and centrifuged at 8000 x g for 10 minutes. The resulting supernatant was transferred to a fresh tube and GSH content was estimated following the Kit’s instructions.

#### Estimation of IC50 to paclitaxel, DOX and cisplatin

To determine the 50% inhibitory concentration (IC50) of PTX, DOX and CIS, HCC1806 and HCC1806R cells respectively were seeded at 6K cells per well in a 96-well plate and cultured in RPMI1640 supplemented with 10% FBS for 24 hours at 37°C with 5% CO2. Following the initial culture period, the media in wells was replaced with fresh media containing one chemo drug with a range of concentration: PTX (0 to 500 nM), DOX (0 to 50 μM), or CIS (0 to 100 μM), followed by incubation for an additional 72 hours; for each drug, duplicate wells were used for each concentration. To assess cell viability, the cells were washed twice with PBS and subsequently incubated for 2 hours with fresh culture media containing 10 μL of CCK-8 solution (catalog no: NC9261855, Fisher Scientific) added in each well. The percentage of viable cells at each concentration was estimated by the absorbance at 450 nm using a Microplate Reader (SpectraMax M5, Molecular Devices). IC50 value for each chemo drug was determined using GraphPad Prism (version 10.0.2).

#### Estimation of cell and tissue cysteine concentration by LC-MASS

5M cells were seeded in a cell culture dish and allowed to culture overnight in culture media (RPMI1640+10% FBS) at 37°C with 5% CO2. Subsequently, the media was replaced with culture media containing CB839 (1 μM), ERA (3 μM), DOX (0.2 μM) and CIS (5μM) and incubated for 24hrs at 37°C with 5% CO2. Following a 24hrs incubation, the cells underwent a wash two times with PBS, and 1ml of extraction solution (40% Methanol, 40% Acetonitrile, 20% DI water, 100mM Formic acid, and 1mM EDTA) was introduced to the cells (26). The extracted cell and solution were gathered using a cell scraper and transferred to tubes on ice. For tissue samples, the tumor tissue homogenization was conducted with the extraction solution mentioned above at 4°C using a Precellys Evolution Homogenizer, which was equipped with Cryolys® Evolution. The 1ml of extraction solution was used to extract 0.1g of tumor tissue. The cell or tissue samples were centrifuged at 16,000g for 10 minutes, and the resulting supernatant was collected. The supernatants were then transferred to tubes on ice. To each tube, 50 μL of internal standard solution (U-^13^C-^15^N-cysteine, catalog no: CNLM-3871-H-PK, Cambridge Isotope Laboratories catalog) was added, followed by the transfer of 450 μL of extracts from the sample tubes to corresponding reaction tubes. Subsequently, 50 μL of triethylamine was added to each reaction tube. To each reaction tube, 5 μL of benzyl chloroformate was added, capped, briefly vortexed, and incubated at 37°C for 10 minutes. After the incubation, the reaction tubes were centrifuged at 6,000g and 4°C for 5 minutes. The supernatant was collected for LC/MS analysis of cysteine. LC/MS (Liquid Chromatography-Mass Spectrometry) data was obtained using a Waters Acquity UPLC system (equipped with a Waters TUV detector at 254 nm and a Waters SQD single quadrupole mass analyzer with electrospray ionization). The LC gradient used was 500 uL/min with a 30-second hold at 95:5 (water: acetonitrile with 0.1% v/v formic acid), a 2-minute gradient to 5:95, and a 30-second hold. An Acquity UPLC HSS C18, 1.7um, 2.1x 50 mm column was employed for the analysis. The obtained data was analyzed using NOVA LC/MS software by Mestrelab Research (https://mestrelab.com/).

#### 4-(3-[^18^F]fluoropropyl)-L-glutamic acid ([^18^F]FSPG) uptake by cells

HCC1806 cells (25,000 cells per well) were seeded in a 96-well strip plate and incubated in complete culture medium supplemented with 10% FBS overnight. Subsequently, the culture medium was exchanged with fresh cell culture medium containing specific drugs including CB839 (1 μM), ERA(3 μM), IKE (3 μM) or DOX (0.2 μM) and cells were incubated for 4hrs. At the end of incubation time, the media were washed twice with PBS and replaced by [^18^F]FSPG (300,000 cpm/well) prepared in PBS containing 0.1% BSA, 10 μM Glutamine, 1 μM Glutamate and 1 μM Cystine with specified concentration of CB839 (1 μM), ERA (3 μM), IKE (3 μM) or DOX (0.2 μM). After incubating for 30min and the supernatant was aspirated, and the wells were washed twice with PBS. After washing, the activity in each well was counted on a γ-counter (2470 Wizard2; Perkin Elmer); afterwards, the amount of protein in each well was estimated by the Lowry method (27). Percent Cell Uptake of [^18^F]FSPG in each well was estimated by the activity in the well normalized to the input activity and protein content.

### Single cell RNA sequencing (scRNAseq)

HCC1806 Cells were cultured in RPMI1640 medium with 10% FBS containing CB839 (1μM), ERA (3μM), or DOX (0.2 μM) for 24 hours. Afterwards, cells were collected by trypsinization and washed 2 times with PBS. Cells were suspended in 0.04%PBS with concentration ≥ 200K cells/100μL for submission to the Genomic and Sequencing Core at the University of Pennsylvania. Next-generation sequencing libraries were prepared using the 10x Genomics Chromium Single Cell 3’ Reagent kit v3 per manufacturer’s instructions. Libraries are uniquely indexed using the Chromium dual Index Kit, pooled, and sequenced on an Illumina NovaSeq 6000 sequencer in a paired-end, dual indexing run. Sequencing for each library was targeted at 20,000 reads per cell. Data is then processed using the Cell Ranger pipeline (10x Genomics, v.6.1.2) for demultiplexing and alignment of sequencing reads to the mm10 transcriptome and creation of feature-barcode matrices.

### scRNAseq data analysis

Using the Cell Ranger (10X Genomics) output of filtered matrices supplied by the sequencing core, the Seurat package (v4) is used to integrate the data for each sample, followed by SCTransform (v2) to perform normalization and unsupervised clustering. In Bioconductor (v3.16), the SingleR package combined celldex databases is used to perform automated cell-type assignments. Cirrocumulus was used for interactive exploration and visualization of datasets. Heatmaps and violin plots were generated using Cirrocumulus. Target genes were extracted from literature (28-31).

### In vivo studies

All animal procedures were approved by the institutional animal care and usage committee (IACUC) of the University of Pennsylvania.

#### Tumor model, treatment, and tumor growth

To establish the human breast cancer xenografts, one million HCC1806 or HCC1806R cells in 100 μL PBS, were inoculated subcutaneously into the right flank of athymic nu/nu mice (female 7-week-old, Charles River). Tumor size was measured by caliber in two orthogonal directions *a* and *b* with *b* being the shorter dimension using formula: V= πab^2^/6.

When the tumor size reached 166 ± 84 mm^3^, the mice were randomly enrolled into one of the 8 groups for treatment for 14 days: Control (no treatment); CB839: 200 mg/kg administered orally twice daily at Day 0, 1, 2, 3, 4, 7, 8, 9, 10, 11, and 14; ERA: 5 mg/kg administered intraperitoneally (i.p.) at Day 0, 2, 4, 7, 9, 11, and 14, CIS: 2.5 mg/kg i.p. at Day 0, 2, 4, 7, 9, 11, and 14; CB839+ERA; CB839+CIS; ERA+CIS; CB839+ERA+CIS. For a combined treatment involving two or three drugs, the dose regime and route were the same as those used for the single-agent treatments. The mouse was sacrificed when the tumor reached the size of 1000 mm^3^ or after being treated for two weeks, whichever occurred earlier, and the tumor was harvested for further analysis.

#### 4-(3-[^18^F]fluoropropyl)-L-glutamic acid ([^18^F]FSPG) PET imaging studies

[^18^F]FSPG was produced at the University of Pennsylvania PET Center. The radiochemical purity of preparations was > 90% and specific activity > 250.1 ± 71.8 mCi/μmol.

Mice bearing HCC1806 xenograft were enrolled when the tumor size reached about 200 mm^3^ by caliber measurement. In vivo PET and CT were performed on the Molecubes modular system (Molecubes Corporation) before and after 2 days of IKE (30 mg/kg per day, i.p.) or PBS (vehicle) treatment. Approximately 7,400 -9,250 kBq (200–250 mCi) of [^18^F]FSPG in 0.2 mL was injected into the tail vein catheter, and a 60-min dynamic PET scan was started immediately and followed by a 2-min CT scan. The dynamic PET data were reconstructed with a temporal resolution of 10 s/frame x 6 frames, 1 min/frame x 9 frames, and 5 min/frame x 10 frames.

Dynamic PET data were reconstructed with a temporal resolution of 10 s/frame × 6 frames, 1 min/frame × 9 frames, and 5 min/frame × 7 frames, and analyzed using PMOD software (version 3.711). A spheric region of interest (ROI) equal to 1/8^th^ of the tumor volume was placed over the tumor region with maximal uptake while avoiding activities from nearby bone structures by referencing to CT images. Similarly, a 2 × 2 × 2 mm cubic ROI was placed over the left ventricle of the heart noting that tracer accumulation in myocardium did not exceed blood pool background. The PMOD software automatically identified the hottest five pixels in the tumor and heart ROI and their average values were used to calculate the time activity curve (TACs) of the tumor and blood, respectively. The ratio of tumor to blood (T/B) signal was estimated from the last frame.

## Statistical Analysis

Data were presented as mean ± standard deviation with sample size specified in figure captions. Statistical analyses were performed in Prism GraphPad (San Diego, CA) with the level of α set at 0.05 for evaluation of significance.

## Data Availability Statement

All data were generated by the authors and are included in the article and its supplementary data files. Single cell RNA sequencing raw data will be deposited in Gene Expression Omnibus (GEO). Raw PET imaging data are available upon request.

## Results

To elucidate the impact of GLSi, xCTi and /or chemotherapy to induce oxidative stress in TNBC cells, we have measured the change of cellular superoxide, the original reactive oxygen species in chemo-naïve HCC1806 cells (**Figure**1a): CB839, ERA, DOX alone or their combination led to a significant increase of the cellular superoxide (*** *P* < 0.001, two tailed t-test for each treatment compared to the control). Notably, <u>dual</u> metabolic blockade (CB839+ERA) induced a significantly higher superoxide level compared to single treatment (*** *P* < 0.001) and the highest superoxide when combined with DOX (*** *P* < 0.001 compared to DOX or dual metabolic blockade). To test whether the increased cellular ROS level leads to apoptosis, we estimated the fraction of cells undergoing early apoptosis by sorting cells double stained with Annexin-V and TO-PRO-3 (**Figure 1b**). Despite increased ROS level, none of the single treatment increased the Q3 (apoptosis) fraction compared to the control (**SI** **Figure 2b**,**c**,**d**), nor did the dual metabolic blockade (**SI** **Figure 2e**) or other 2-drug combinations (**SI** **Figure 2f**,**g**) despite significantly increased ROS level. Combination of DOX with dual metabolic blockade led to a significant fraction of apoptotic cells (**Figure 1b**). Data summarized in **Figure 1c** confirmed this binary behavior, which suggests the existence of a threshold of oxidative stress, under which cancer cells can deploy antioxidative mechanisms to resolve the stress and survive, whereas over which these mechanisms are depleted, and massive cell death ensues.

To evaluate whether this approach facilitates overcoming chemo-resistant TNBC, we employed a paclitaxel-resistant version of HCC1806 line, referred to as HCC1806R that has 35-, 4- and 3-fold increase of IC50 to PTX, DOX, and CIS, respectively (**Figure 2a**). When the parent and resistant cells were exposed to the <u>same</u> drug combination / concentration/ exposure time (24 h), the resistant cells produced significantly lower superoxide levels across all single or combined treatment compared to the parent cells (**Figure 2b**). These data suggest that resistant cells have increased capacity of mitigating oxidative stress, however, dual metabolic blockade elevated superoxide level significantly (**** *P* < 0.0001 compared to single agent, **Figure 1d**), confirming its effectiveness in inducing oxidative stress in resistant TNBC cells. Similar to the parent cells, significantly increased apoptosis was induced by dual metabolic blockade combined with CIS chemotherapy in HCC1806R cells (**Figure 1e**).

**Figure 2.**
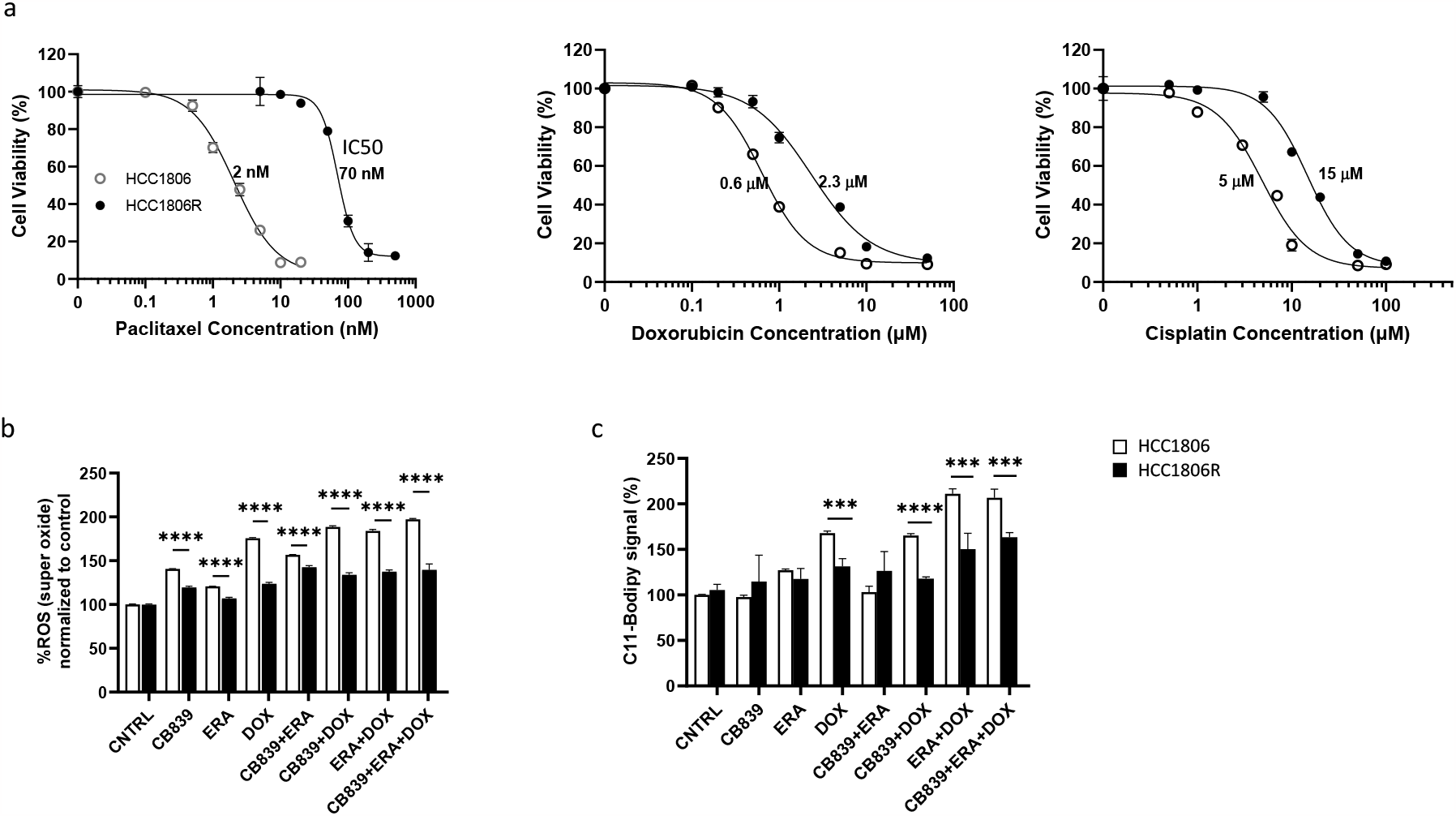
Differential IC50 to chemotherapy drugs, levels of superoxide and ferroptosis in parent *versus* resistant cells induced by dual metabolic blockade and chemotherapy drugs. **a**: IC50 of HCC1806 and its resistant counterpart (HCC1806R) to paclitaxel (PTX), DOX and CIS, respectively. Cellular super oxide (**b**) and ferroptosis level (normalized to controls set at 100%) (**c**) in HCC1806 and HCC1806R cells exposed to CB839, ERA, DOX or their combinations were presented relative to CNTRL. Mean and SD were presented for b and c. *** *P*<0.001, **** *P*<0.0001 comparing parent with resistant cells. IC50 to PTX plot was a reprint from Choi et al (25).

Besides inducing apoptosis, ROS causes lipid peroxidation, which is an irreversible process, leading to cell death via iron-dependent ferroptosis pathway (14). C11-BODIPY, a fluorescent dye molecule that binds to oxidized lipids, is a classic marker for ferroptosis (16). As shown in **Figure 3a**, DOX and ERA induced a robust C11-BODIPY signal whereas CB839 did not. C11-BODIPY signal was significantly reduced in the presence of iron chelator deferoxamine (DFO) (**** *P* < 0.0001 compared to without DFO) or Ferrostatin-1(Fer-1), a known inhibitor of ferroptosis (**** *P* < 0.0001 compared to without Fer-1), confirming the nature of C11-BODIPY signal as a marker of ferroptosis. In parent cells (**Figure 3b**), ERA+DOX elevated the level of ferroptosis over DOX or ERA alone (**** *P*<0.0001). In the <u>resistant</u> cells (**Figure 3c**), while DOX alone was still able to induce higher level of ferroptosis (\*\**P*< 0.01 compared to the CNTRL), ERA+DOX was unable to further elevate ferroptosis level (*P=* 0.3429, compared to DOX alone) hence dual metabolic blockade plus DOX was necessary to increase ferroptosis above what was obtained by DOX (\*\*\**P<* 0.001). Significantly lower C11-BODIPY signal was observed in HCC1806R cells compared to the parent cells in response to DOX alone or DOX+ treatment(s) (**Figure 2c**), confirming the enhanced ability of resistant cells to mitigate ferroptotic cell death induced by oxidative stress.

**Figure 3.**
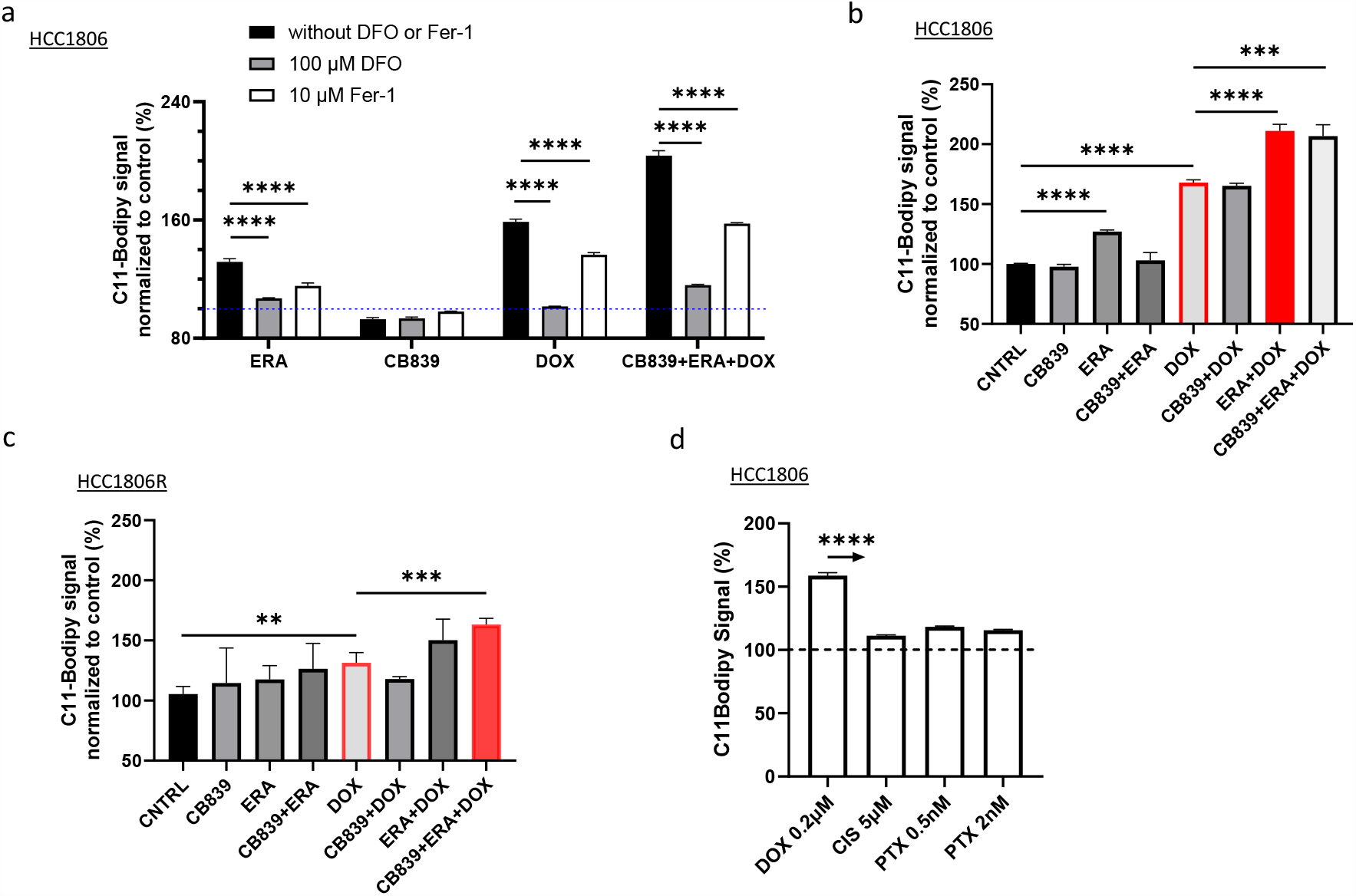
Dual metabolic blockade and chemotherapy drugs mediated ferroptosis in chemo naive and resistant TNBC cells. Ferroptosis was measured by C11-BODIPY signal (normalized to controls set at 100% by dotted blue line). **a:** C11-BODIPY signal in the presence or absence of DFO, or Fer-1. C11-BODIPY signal in HCC1806 (**b**) and HCC1806R (**c**) cells after 24 h exposure to CB839, ERA, DOX or their combinations. **d**: Chemotherapy drugs have different capacity to induce ferroptosis. \*\**P*<0.01, \*\*\**P*<0.001, \*\*\*\**P*<0.0001. Data were presented as mean and SD (N=4 replicates in each group). In panel d, arrow indicates *P* < 0.0001 comparing DOX with other drugs at specified concentration.

Our data revealed different ability of chemotherapy drugs to induce ferroptosis (**Figure 3d**): while DOX induced robust ferroptosis above the control, CIS, even applied at 25-fold higher concentration had minimal impact on ferroptosis, so did PTX when applied at the IC50 concentration (2 nM) or below. In the in vivo study, we chose to use CIS as chemotherapy to avoid a significant contribution of DOX-mediated ferroptosis to the treatment effect. In addition, HCC1806 and HCC1806R are much less sensitive to CIS than DOX or PTX (Figure 2a). Therefore, using CIS allowed us to test the efficacy of dual metabolic inhibition for sensitizing chemo resistant TNBC.

In our *in vivo* xenograft models, while CIS retarded the growth of chemo naïve HCC1806 tumors (**Figure 4a**), it failed to induce growth delay in HCC1806R tumors compared to PBS controls (blue symbols, **Figure 4b**), confirming that these tumors are resistant to CIS. GLSi or xCTi alone did not impact the tumor growth either (**Figure 4c**). CB839 or ERA combined with CIS or CB839+ERA delayed the tumor growth significantly albeit moderately (green symbol, **Figure 4d** P< 0.05 compared to the control or single drug regimen). dual metabolic inhibition combined with CIS induced the most significant tumor growth delay (red symbol, **Figure 4e, \*\*** *P* <0.01 comparing to control, ERA alone, CB839+CIS, or ERA+CIS, details in figure caption).

**Figure 4.**
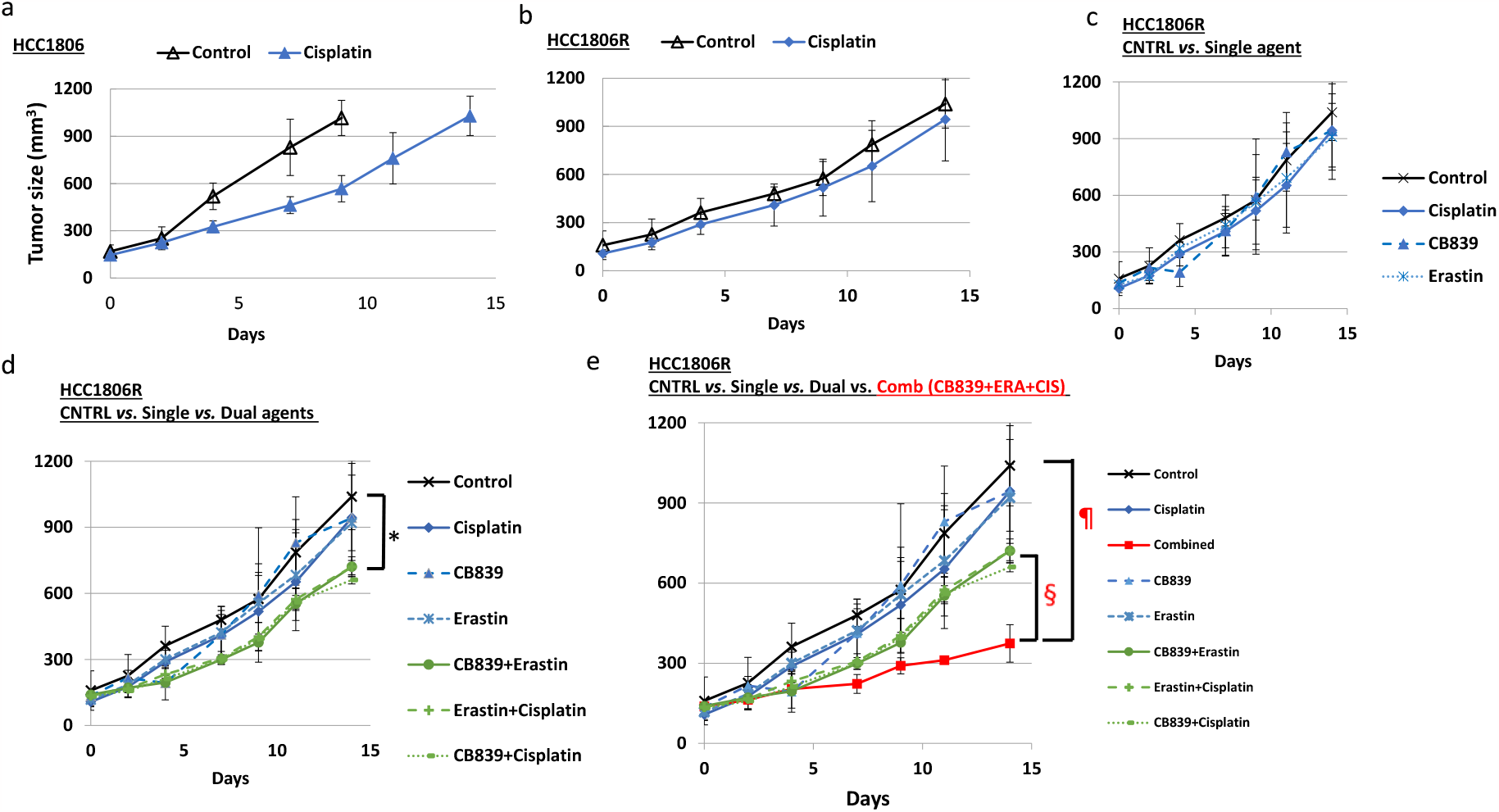
Double metabolic inhibition sensitizes chemo-resistant TNBC to cisplatin chemotherapy. Tumor growth time course of HCC1806R xenografts in response to 2-week treatment by CB839, Erastin, Cisplatin or their combinations: CB839 (200 mg/kg twice daily oral), Erastin (5mg/kg, ip) and CIS (2.5 mg/kg, ip) given 3 times /wk x 2 wks. Comb =CB839+ERA+CIS. ^¶^*P* = 0.002, 0.021, 0.016 and 0.004 comparing Comb vs. CNTRL, Cisplatin, CB839 and Erastin, respectively. ^§^*P* = 0.026, 0.002, 0.003, comparing Comb vs. CB839+ERA, CB839+CIS, and ERA+CIS, respectively; ^*^*P* < 0.05 comparing CB839+CIS, ERA+CIS vs CNTRL.

Superoxide and C11BODIPY signal were elevated upon 24 h exposure to metabolic inhibitors and/or chemotherapy drug, resulting in apoptosis and ferroptosis in TNBC cells (Figure 1 and 3). However, these pharmacological interventions were unable to induce transcriptome changes as revealed by scRNAseq analysis (**Figure 5a**): no change in mRNA level of GLS or SLC7A11 (xCT) after 24 h exposure to CB839, ERA, or DOX compared to CNTRL (heat map and violin plot). ACSL4 (Acyl-CoA Synthetase 4) expression, which regulates ferroptosis (28), had no change, nor did expression of genes regulating ROS scavenging, GSH synthesis, NADPH production and other ROS mitigation mechanisms (19) (**SI Figure 3**). No change was detected in the expression of NRF2, the master transcription factor of redox homeostasis (19) within the 24 h time frame (Figure 5a).

**Figure 5.**
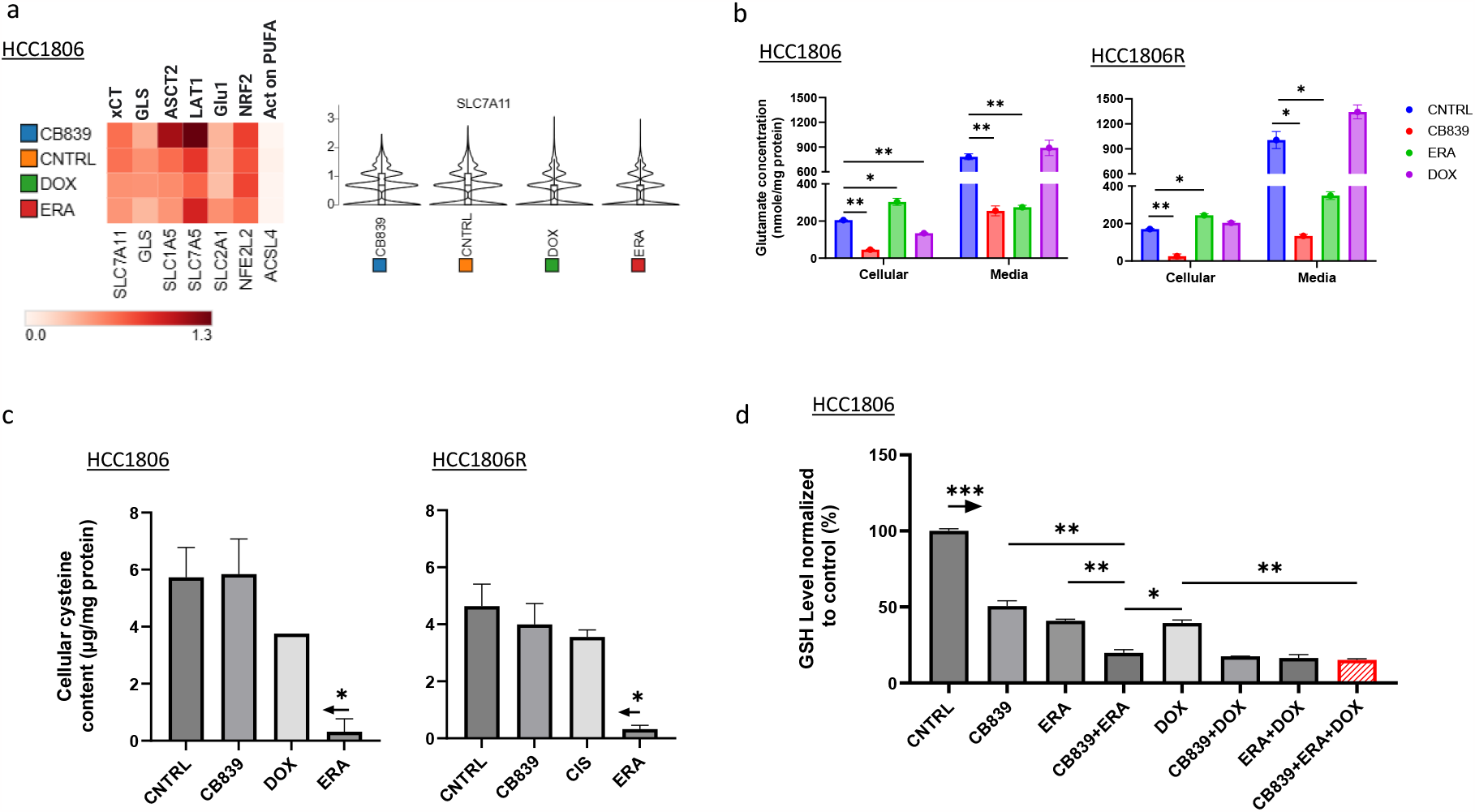
Changes of metabolites precede changes in gene expression (mRNA) in TNBC cells responding to metabolic blockade and/or chemotherapy. **a**: single cell RNA sequencing analysis revealed no changes in the expression of GLS and xCT after 24 h exposure to CB839, ERA, or DOX; violin plot of SLC7A11 (xCT) expression in treated cells. **b:** Intra and extracellular Glu concentration after specified treatment (duplicate samples for each treatment. Drug concentrations are specified in Methods). **c:** Cellular cysteine level in HCC1806 and HCC1806R cells after specified treatment (duplicate samples for each treatment. Drug concentrations are specified in Methods). Arrow above ERA indicates \**P*<0.05 compared to CNTRL, CB839 or DOX treatment, respectively. **d:** Cellular GSH level in HCC1806 cells after 24 h exposure to specified treatment (duplicate samples for each treatment). Drug concentrations are specified in Methods). In d, arrow above the CNTRL indicates *P*<0.001 comparing to each treatment in the panel. HCC1806 or HC C1806R cells were incubated with ERA (3μM), CB839 (1μM), Dox (0.2μM) or their combinations for 24 h before being processed for metabolites and scRNAseq analysis.

In contrast to lack of changes in mRNA level, remarkable changes of intra and extracellular metabolites were captured (**Figure 5b**): after 6h incubation with CB839 in culture media free of Glu and FBS, both intra- and extracellular Glu concentration were significantly reduced compared to CNTRL, consistent with reduced cellular Glu pool due to GLS inhibition. Meanwhile, by blocking xCT antiport activity, ERA increased cellular Glu concentration significantly (** *P* <0.01 compared to CNTRL) due to reduced Glu export. DOX reduced cellular Glu concentration only in parent cells (** *P* <0.01 comparing to CNTRL), suggesting increased Glu consumption for GSH synthesis in response to DOX-induced oxidative stress, whereas such response was blunted in resistant cells. Concentration of cysteine, formed by two cystine molecules after being transported inside the cell, exhibited a significant reduction in both parent and resistant cells in response to ERA, consistent with xCT blockade (*P* <0.05, compared to CNTRL) whereas CB839 or DOX had no effect (**Figure 5c**). Cellular GSH level was decreased sharply after 24 h exposure to CB839, ERA, or DOX (**Figure 5d**); notably, dual metabolic blockade combined with DOX almost depleted cellular GSH and this treatment also induced the highest level of apoptosis (Figure 1c), suggesting a link between GSH depletion and the superoxide threshold over which apoptosis ensues.

The large change of metabolites concentration proceeding changes of gene expression provides a rationale for development biomarkers based on cellular metabolite pool size. We evaluated the utility of [^18^F]FSPG, an analog of glutamate, as a marker for xCTi. Like ERA, IKE is a potent xCTi but it is more metabolically stable and exhibits improved bioavailability *in vivo* (32,33), therefore, IKE was utilized in our PET study: comparing at baseline and post IKE or VEH treatment, [^18^F]FSPG PET signal was reduced notably only after IKE treatment (**Figure 6a**) and the tumor-to-blood ratio (T/B) of [^18^F]FSPG was decreased significantly in IKE treated mice compared to baseline whereas T/B had no change in VEH treated mice (**Figure 6b**). Cell uptake study of [^18^F]FSPG confirmed the *in vivo* results showing highly significant reduction of ^18^F activity in the HCC1806 cells after ERA or IKE treatment (**** *P* <0.0001 compared to CNTRL) as well as CB839 (*** *P* <0.001) whereas DOX appeared to have no effect on FSPG uptake (**Figure 6c**). Note that the cells were incubated for 24 h with CB839, ERA, IKE, or DOX before cell uptake study (Methods). In tumors harvested after PET imaging, a significant reduction of cysteine level was detected in IKE treated group (* P<0.05 compared to VEH, **Figure 6d**).

**Figure 6.**
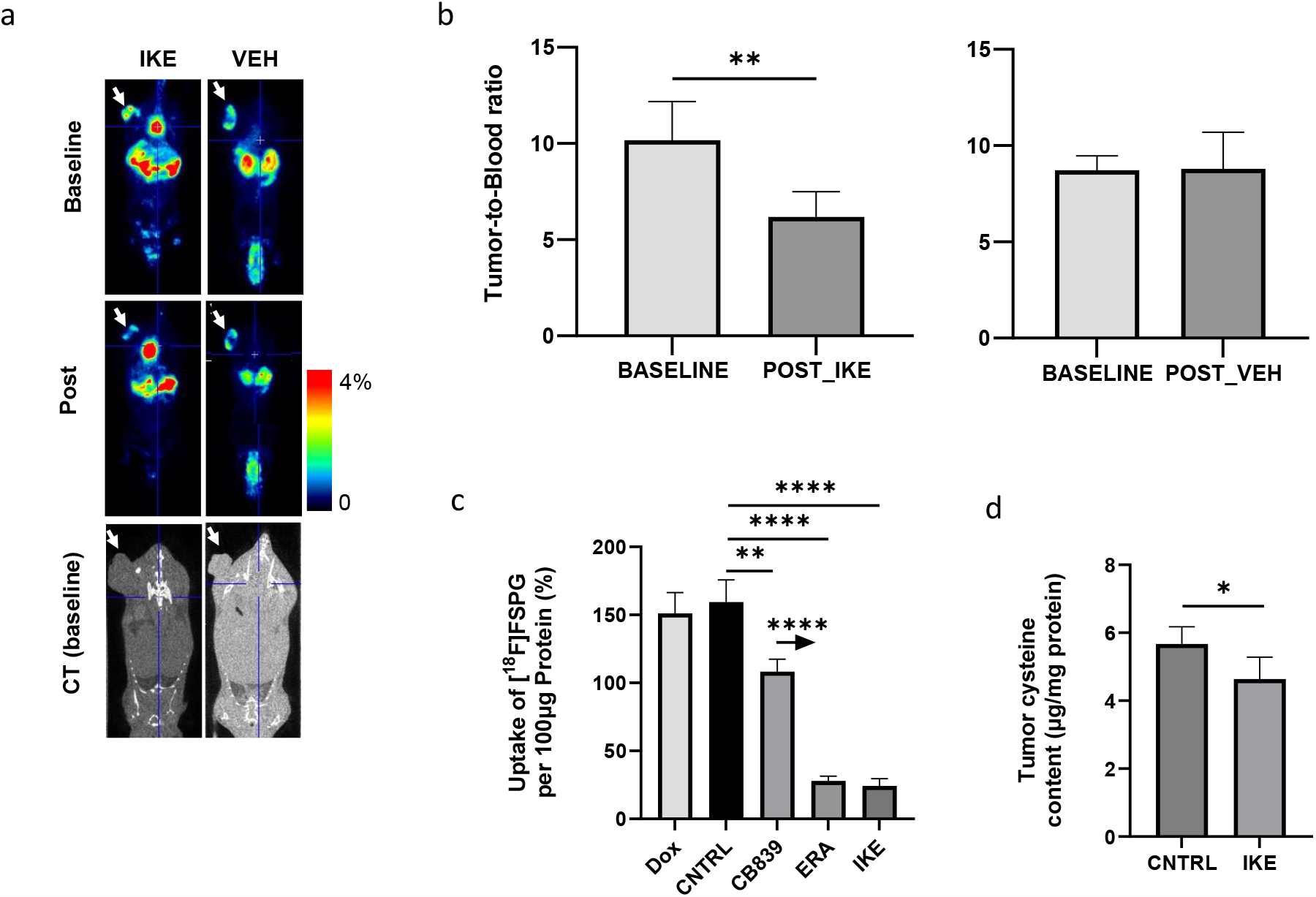
PET imaging marker for xCT blockade. **a**: Representative in vivo [^18^F]FSPG PET and CT images of human TNBC xenograft (HCC1806) at baseline and after xCT inhibitor (IKE) or vehicle (PBS) treatment. Tumor is pointed by the white arrow. **b:** Group average and statistics of tumor-to-blood ratio (T/B) at baseline and post-IKE or VEH. **c:** HCC1806 cell uptake of [^18^F]FSPG after 30 min of incubation with the tracer in PBS solution containing 0.1% BSA, 10 μM Glutamine, 1 μM Glutamate, 1 μM Cystine (N=4 for each treatment). The cells were incubated with specified drug for 24 h before cell uptake study. In c, the arrow above CB839 indicates *P*<0.0001 comparing to ERA or IKE treatment respectively. **d:** Cysteine levels in HCC1806 tumors after IKE or VEH treatment (N=4 for each treatment). IKE dose regimen is specified in Methods.

## Discussions

Enhancing oxidative stress in rapidly dividing cancer cells is an important factor mediating the efficacy of many cytotoxic chemotherapy agents (34-37). Our study revealed that chemo-resistant cells derived from highly glutaminolytic TNBC were able to maintain lower levels of cellular superoxide and lipid peroxidation (ferroptosis) in the face of oxidative challenge compared to their chemo-naïve counterparts (Figure 2b-c), providing a plausible mechanism for resistance and a motivation for targeting glutamate production and export to abrogate chemotherapy resistance. Our prior work using both radioisotope and stable isotope of glutamine revealed that glutamine metabolism through GLS contributing primarily to a large pool of cellular glutamate (38,39) that could be reserved for mitigating oxidative stress (SI Figure 1) besides feeding the TCA cycle. Leveraging this insight, we investigated the potential of disrupting redox balance through dual metabolic blockade as a strategy to sensitize chemo resistant TNBC tumors to CIS and DOX chemotherapy.

Dual metabolic blockade led to 80% depletion of GSH (Figure 5d) as the result of reduced cellular Glu and depleted cysteine (Figure 5b and c). GSH depletion prevents the formation of CIS-GSH conjugates, which are exported out of cells thereby allowing cancer cells to evade CIS-mediated DNA damage and cell death (40), therefore, our data supports GSH depletion as mechanism underpinning the sensitization of resistant TNBC to CIS chemotherapy: CIS plus dual metabolic blockade culminated in the highest levels of apoptosis in resistant cells (Figure 1e) and the most significant growth delay in resistant tumors (Figure 4e). The dual metabolic blockade when combined with DOX also led to highly significant apoptosis (Figure 1c) likely because the high superoxide level induced by such combination (Figure 1a) was not mitigated by antioxidative mechanisms, triggering apoptotic death. Meanwhile, DOX’s ability to induce additional cell death in the form of ferroptosis was bolstered significantly by addition of ERA (red bar in Figure 3b). Overall, <u>dual</u> metabolic blockade plus DOX facilitates overcoming the chemo resistance by significantly enhancing both apoptosis and ferroptosis over those obtained by DOX alone (Figure 1c and Figure 3b-c).

Our data suggest the existence of a threshold for superoxide to trigger apoptosis, which sheds an insight into the lack of cytotoxic effect of CB839 as single agent: although it robustly increased cellular super oxide level (Figure 1a) that consequently diminished cellular GSH pool by 50% (Figure 5d), CB839 had no impact on inducing apoptotic (Figure 1c,e) or ferroptotic cell death (Figure 3a-c). In vivo, CB839 did not enhance CIS chemotherapy in resistant tumors (P = 0.132 comparing CB839+CIS vs. CIS, Figure 4d), mirroring clinical trials outcome.

Our results reveled significant changes in cellular metabolites (Glu, GSH and cysteine) preceding transcriptome changes of targeted proteins (Figure 5b,c,d). We demonstrated the utility of [^18^F]FSPG, which is a glutamate analog (41) and has been evaluated in clinical studies (42-44), for detecting xCT blockade by ERA or IKE as early as 24 hours in cells or 2 days of treatment in mice (Figure 6). Our data are consistent with reports from McCormick et al (45) and Greenwood et al (46) that show the [^18^F]FSPG retention in cells correlating with cellular cystine level and coinciding with markers of oxidative stress or antioxidant capacity. These findings underscore the potential of developing non-metabolized amino acid PET approaches to capture early metabolic changes such as those induced by GLSi or glutamine transporter inhibitor (27,47-50). In summary, our study provides compelling evidence for the therapeutic benefit and feasibility of dual metabolic blockade as a translational strategy to sensitize resistant TNBC to cytotoxic chemotherapy.

## Acknowledgement

We thank Calithera Biosciences Inc. and Dr. Marina Gelman for supplying CB839. This study was supported by R01CA233771, R01CA211337 and R01CA266285, and by a grant from Ludwig Institute for Cancer Research. CH was supported by training grant (5T32EB004311) and an RSNA Resident Research grant (RR2121).

