## Supplemental Figures for "Disruption of redox balance in glutaminolytic triple negative breast cancer by inhibition of glutamate export and glutaminase"

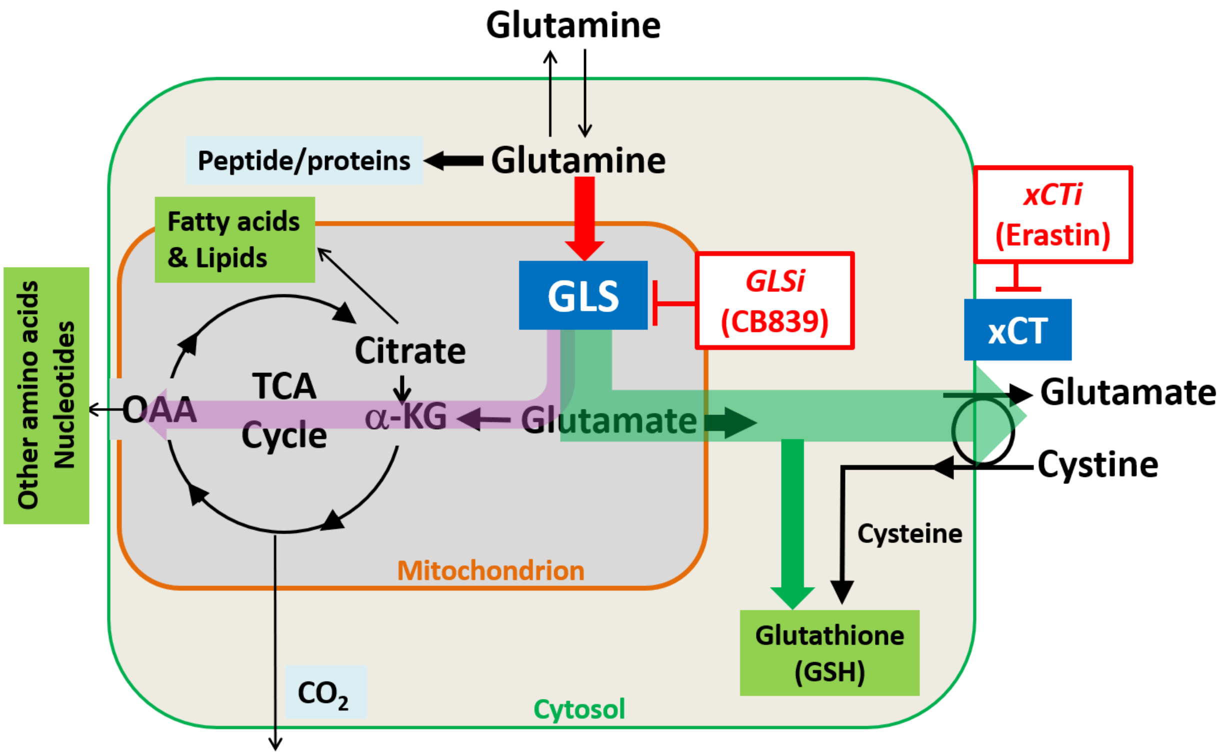


**SI Figure 1 Schematic diagram describing rationales of dual metabolic inhibition by GLSi (e.g., CB839) and xCTi (e.g., Erastin).** Following entry of glutamine into the cell, the purple arrow is the glutaminolysis pathway while the green arrows indicate glutamate contributions to de novo glutathione pathway. *xCT* (SLC7A11): glutamate/cystine antiporter which is inhibited by Erastin; GLS: Glutaminase, which is specifically inhibited by CB839, an investigational *GLSi* in phase-2 trial.

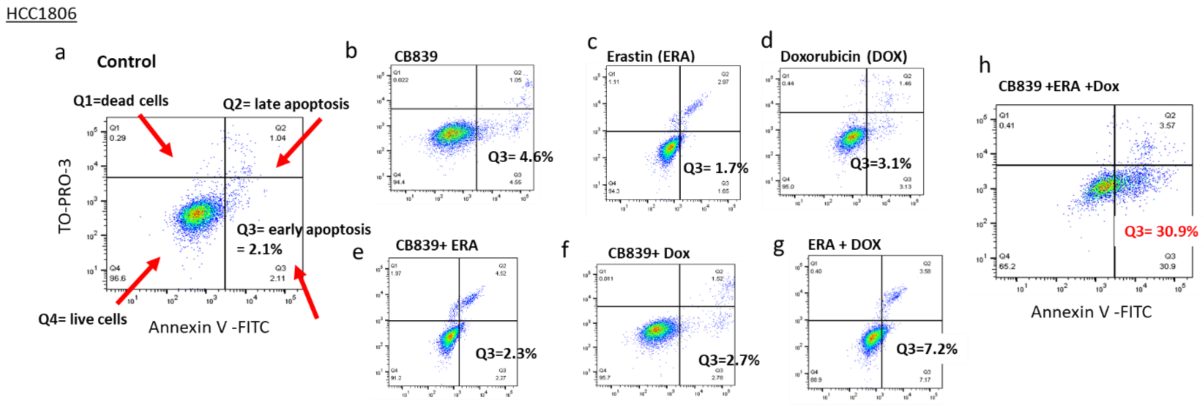


**SI Figure 2 FACS based estimation of early apoptosis in TNBC cells induced by CB839, ERA, doxorubicin (DOX) or their combinations.** Fraction of cells undergoing apoptosis (high Annexin V-FITC but low TO-PORO-3 signal) are in the 3rd quarter (Q3).

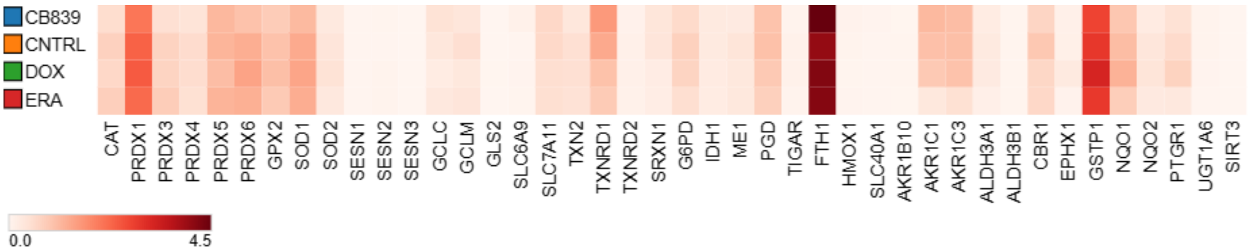


**SI Figure 3 Expression of genes controlled by NRF2 transcription factor and involved in maintaining redox homeostasis in cancer cells** (reference to Hayes et al (19)).
